# Protection of the Telomeric Junction by the Shelterin Complex

**DOI:** 10.1101/2024.08.18.608453

**Authors:** Sajad Shiekh, Darion Feldt, Amanda Jack, Sineth G. Kodikara, Janan Alfehaid, Sabaha Pasha, Ahmet Yildiz, Hamza Balci

## Abstract

Shelterin serves critical roles in suppressing superfluous DNA damage repair pathways on telomeres. The junction between double-stranded telomeric tracts (dsTEL) and single-stranded telomeric overhang (ssTEL) is the most accessible region of the telomeric DNA. The shelterin complex contains dsTEL and ssTEL binding proteins and can protect this junction by bridging between the ssTEL and dsTEL tracts. To test this possibility, we monitored shelterin binding to telomeric DNA substrates with varying ssTEL and dsTEL lengths and quantified its impact on telomere accessibility using single-molecule fluorescence microscopy methods *in vitro*. We identified the first dsTEL repeat nearest to the junction as the preferred binding site for creating the shelterin bridge. Shelterin requires at least two ssTEL repeats while the POT1 subunit of shelterin that binds to ssTEL requires longer ssTEL tracts for stable binding to telomeres and effective protection of the junction region. The ability of POT1 to protect the junction is significantly enhanced by the 5’-phosphate at the junction. Collectively, our results show that shelterin enhances the binding stability of POT1 to ssTEL and provides more effective protection compared to POT1 alone by bridging single- and double-stranded telomeric tracts.

**Table of Content Graphic:** 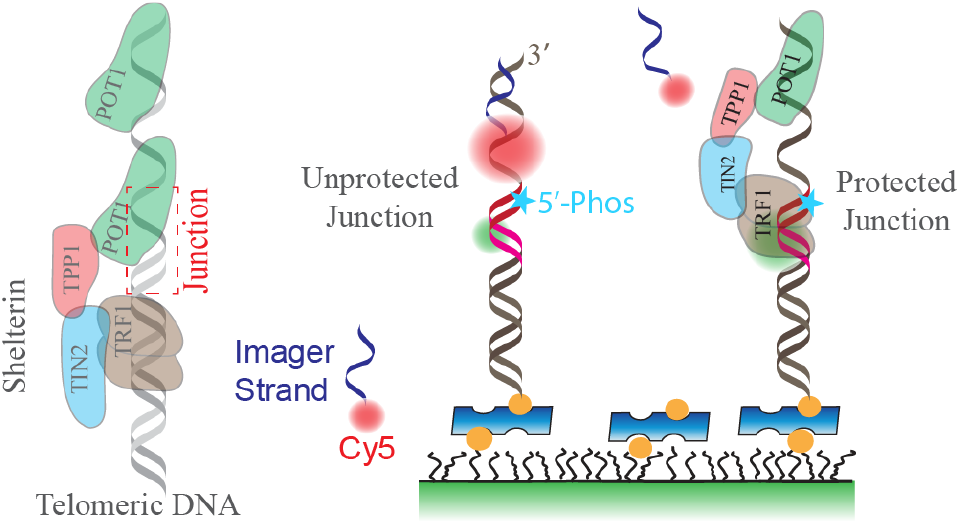

## Introduction

Telomeric DNA serves critical functions in protecting and maintaining the ends of linear chromosomes of eukaryotes, which are prone to end-protection and end-replication problems ^1,2^. The repeating TTAGGG (called a G-Tract) sequence of telomeres has unique characteristics that enable it to form the t-loop and G-quadruplex (GQ or G4) structures to remedy these problems ^3,4^. In addition, telomeres recruit a six-protein shelterin complex, which plays critical roles in distinguishing telomeric ends from DNA double strand breaks ^5–7^. TRF1 and TRF2 subunits of shelterin bind to dsTEL ^8,9^, while POT1 binds to ssTEL ^10^. RAP1 forms a complex with TRF2 to regulate its specificity to telomeres ^11^, while TIN2 and TPP1 interact with TRF2, TRF1, and POT1 to establish a protein-mediated bridge between ssTEL and dsTEL ^12,13^. Deletion or dysfunction of shelterin proteins activates various DNA damage repair pathways at telomeres ^5,7^.

Protection of telomeric overhangs by POT1 or shelterin complex has been studied *in vitro* and *in vivo* ^5,14–17^. Ensemble ^18,19^ and single molecule ^20–23^ studies have revealed important insights about the folding patterns and compactness of the telomeric overhangs. In particular, single molecule fluorescence studies have identified the junction region to be the most accessible region in the overhang in the absence of shelterin ^20,23^. Consistent with this finding, recent structural studies demonstrate recognition of this junction by human POT1 through the phosphorylated 5’-end of chromosome ^24^. These studies show the capping of this junction by a particular surface of POT1 (called the “POT-hole” surface) and mutations in this surface weaken junction protection and compromise suppression of DNA damage response ^24^. While shelterin has been shown to provide significant protection throughout the overhang ^24,25^, exclusion of TIN2 and TPP1 that mediate the connection between POT1 and TRF1/TRF2 or removing the dsTEL segment from the DNA constructs eliminated the elevated protection provided by shelterin compared to the POT1-only case ^25^. These studies hinted that shelterin components may form a protein bridge between ssTEL and dsTEL at the junction, but this possibility has not been carefully tested in minimal telomeric constructs that contain the junction region.

In this study, we focused on the junction region between ssTEL and dsTEL by performing single molecule Förster Resonance Energy Transfer (FRET) ^26^, FRET-Point Accumulation for Imaging in Nanoscale Topography (FRET-PAINT)^27,28^, and Protein-Induced Fluorescence Enhancement (PIFE) experiments on DNA constructs that have 1-2 telomeric dsTEL repeats and 1-3 ssTEL repeats^29^. These experiments revealed that shelterin reduces the accessibility of the telomeric junction by simultaneously interacting with the ssTEL and dsTEL tracts and provided more efficient sequestration than POT1 alone.

## Results and Discussion

We performed measurements on DNA constructs with the C-rich strand terminating with the AATCCC-5’ or CCAATC-5’ sequence (named 5’-CCC and 5’-CTA constructs, respectively), which is identified to be the terminal sequence in majority of human cells^30^. We also introduced phosphate to the 5’-end of the C-rich strand to investigate its impact on protection by POT1 and shelterin.

### The shelterin binding site on dsTEL

To enable shelterin to establish a protein bridge across the telomeric junction, we designed short telomeric constructs that contain two dsTEL repeats and 1-3 ssTEL G-Tracts. The C-rich strand ended with the AATCCC-5’ sequence in these constructs (5’-CCC constructs, Table S1). The overhang of these constructs cannot form intramolecular G-quadruplex (GQ) structures because GQ formation requires at least four G-Tracts. In addition, these constructs cannot form intermolecular GQs because DNA concentrations during annealing and single molecule imaging were kept several orders of magnitude lower than those required for intermolecular GQ formation.

To identify whether shelterin prefers to bind the first or second dsTEL repeat from the ssTEL/dsTEL junction, we performed PIFE experiments on DNA constructs labeled with a Cy5 dye at the 4^th^ bp within the first telomeric repeat or the 10^th^ bp within the second telomeric repeat. The enhanced fluorescence observed in PIFE assays is attributed to reduction in photoisomerization rate of the fluorophore from the brighter (trans) to the dimmer (twisted or cis) state upon interaction with a proximal protein ^31,32^. As this effect is not restricted to proteins but observed for other biomolecules as well, “photoisomerization-related fluorescence enhancement” has been proposed as a broader and more accurate name for the effect, which maintains the PIFE acronym ^33^. We directly excited the Cy5-fluorophore and quantified PIFE signal before and after introducing the 4-component human shelterin complex ^34^ (TRF1-TIN2-TPP1-POT1; shelterin hereafter) (Fig. 1). We observed an approximately 2-fold increase in fluorescence intensity in constructs with Cy5 at the 4^th^ bp from the junction, whereas the intensity remained unchanged in constructs with Cy5 at the 10^th^ bp, regardless of the number of repeats in ssTEL. This suggests the first dsTEL repeat, and not the second repeat, from the junction is the preferred site for establishing the shelterin bridge for the 5’-CCC constructs. Measurements performed on the same DNA constructs in the presence of POT1-only did not show a significant PIFE signal, suggesting the fluorescence enhancement is specifically due to the establishment of the shelterin bridge across the junction.

**Figure 1.**
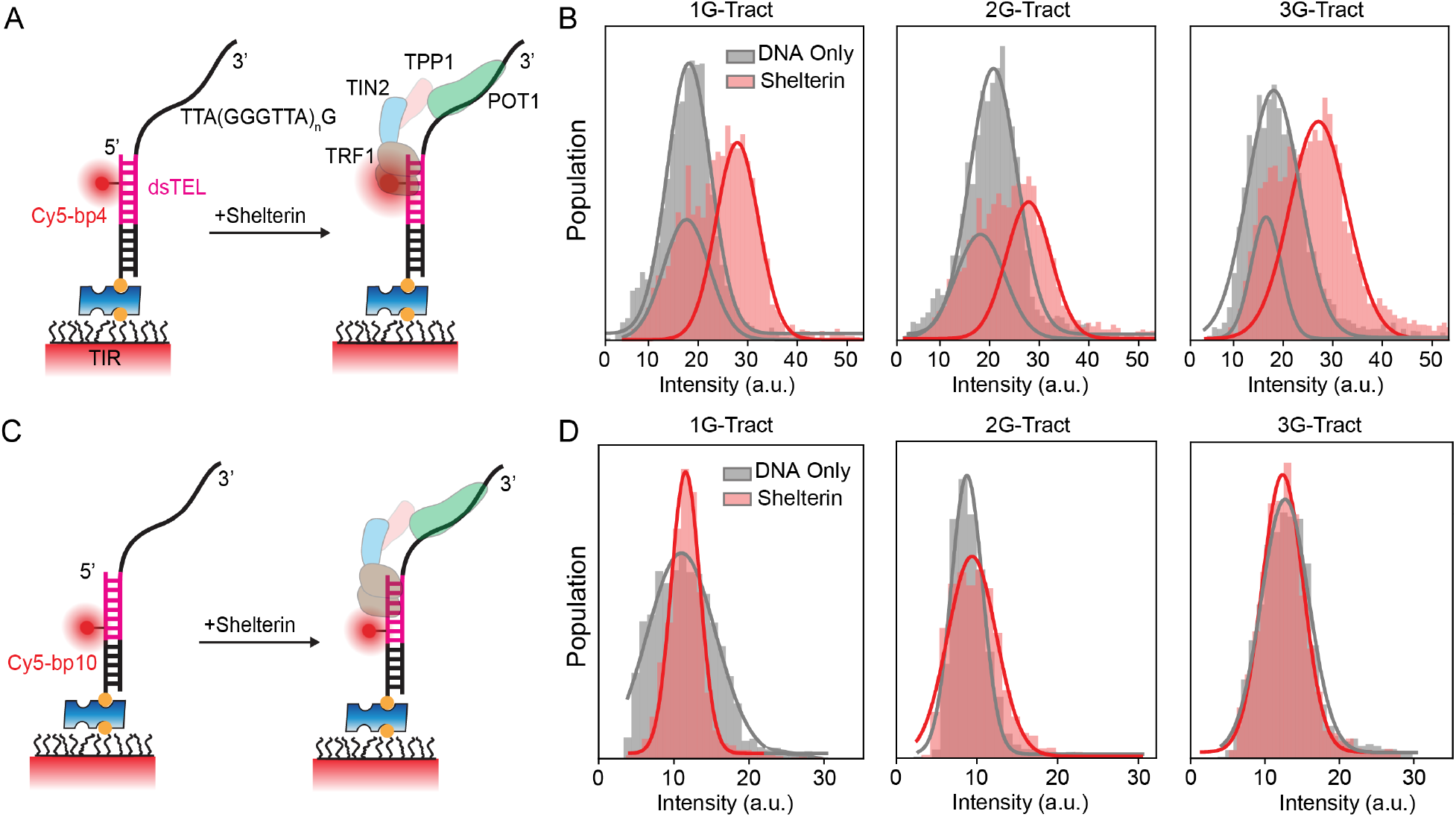
Shelterin binds to the dsTEL nearest to the junction. (A) Schematics of the 5’-CCC DNA constructs in which the C-rich strand terminates with TAACCC-5’ sequence and Cy5 fluorophore is placed in the 4^th^ bp from the ssTEL/dsTEL junction (within the first telomeric repeat in dsTEL). The ssTEL (overhang) contains *n* = 1, 2, or 3 G-Tracts. (B) Normalized population vs. Cy5 intensity on the DNA constructs shown in (A) before and after they interact with shelterin. Shelterin addition resulted in a PIFE signal (red Gaussian peak) in all constructs suggesting that shelterin interacts with the telomeric repeat closest to the junction on dsTEL. (C) Schematics of similar 5’-CCC DNA constructs in which the Cy5 is placed in the 10^th^ bp from the ssTEL/dsTEL junction (within the second telomeric repeat in dsTEL). (D) PIFE experiments on constructs shown in (C) before and after they interact with shelterin. None of the constructs show significant PIFE, suggesting shelterin preferentially interacts with the first repeat on dsTEL rather than the second.

### Minimum ssTEL length for shelterin binding

Next, we sought to determine the minimum ssTEL length required for establishing a shelterin bridge across the junction for the 5’-CCC constructs. We kept two telomeric repeats in dsTEL and varied the ssTEL length between 1-3 G-Tracts. The DNA constructs were labeled with Cy5 at the 4^th^ bp of dsTEL and with Cy3 at the 3’-end of the overhang for smFRET measurements in the presence of shelterin or the POT1 subunit of shelterin (Fig. 2). POT1 contains two DNA binding domains (DBDs) that bind to 9 nt long telomeric sequence (TAGGGTTAG)^10^. The 1G-Tract (10 nt long ssTEL) and 3G-Tract (22 nt long ssTEL) constructs can accommodate stable binding of one and two POT1 molecules, respectively, whereas the 2G-Tract (16 nt long ssTEL) can accommodate both DBDs of one POT1 and only one of the DBDs of the second POT1.

**Figure 2.**
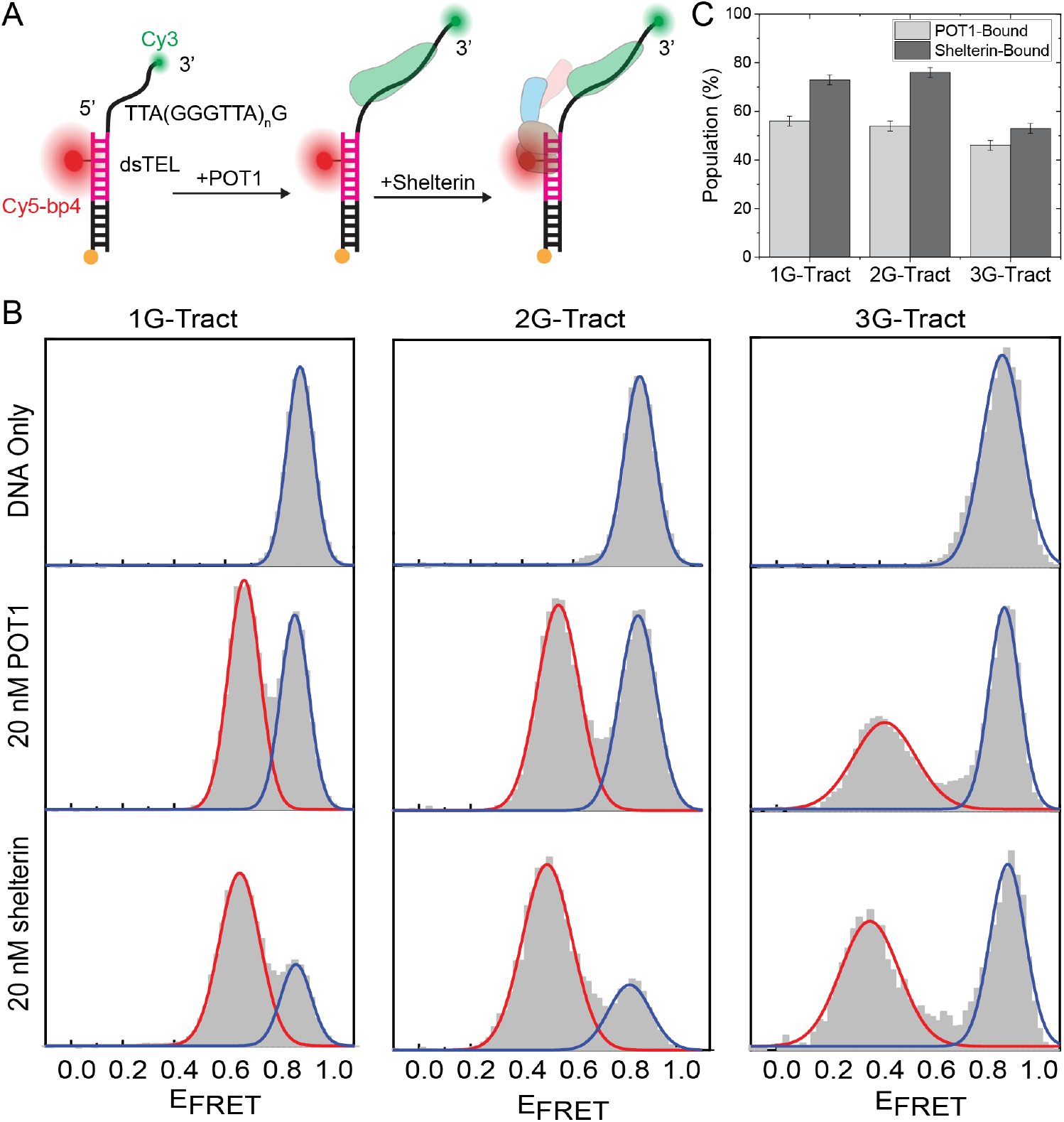
Shelterin more efficiently binds to the telomeric junction than POT1 alone. (A) Schematics of 5’-CCC DNA constructs with 1-3 G-Tracts in ssTEL, 2-repeats in dsTEL, and Cy5 in the 4^th^ bp from the junction. The binding of POT1 or shelterin to the DNA constructs results in the emergence of a peak at a lower FRET efficiency than the DNA-only case. (B) Normalized population vs. FRET efficiency histograms for DNA constructs with 1, 2, or 3 G-Tracts in ssTEL. Solid curves represent a fit to a function with multiple Gaussian peaks. Blue and red curves represent DNA-only and protein-bound populations, respectively. Number of molecules for each measurement and details of Gaussian fits are given in Table S2 and Table S3. (C) Percent population of POT1-bound and Shelterin-bound states based on the areas under the respective peaks. Error bars were estimated based on the fitting errors. T-test analysis showed the variation among POT1 and shelterin cases to be statistically significant (p<0.001 for 1G-Tract and 2G-Tract and p=0.005 for 3G-Tract, Table S4).

Due to the flexibility of the ssTEL, we detected a high FRET signal (E_FRET_ = 0.89 – 0.91; DNA only population) between Cy3-Cy5 FRET pairs for all DNA constructs in the absence of proteins. The binding of POT1 to the 1G-Tract construct resulted in a reduction in the FRET signal (E_FRET_ = 0.67 ± 0.02) due to the stretching of the ssTEL. The FRET efficiency (E_FRET_) of the peak representing the POT1-bound population progressively reduced from 0.67 to 0.42 as the ssTEL length is increased from one to three G-Tracts. Similarly, the addition of shelterin resulted in E_FRET_ to shift from 0.66 to 0.36 from the 1G-to 3G-Tract constructs (Tables S2 and S3).

The populations of the peaks observed in smFRET measurements provided further insights about POT1 binding. The low E_FRET_ population that emerges upon shelterin binding constitutes a greater fraction of the total population compared to that for the POT1-only case (Fig. 2C). The populations of POT1 and shelterin cases are 56% and 73% for 1G-Tract, 54% and 76% for 2G-Tract, 46% and 53% for 3G-Tract constructs, respectively (Fig. 2C and Table S3). These results are consistent with TPP1 enhancing the binding affinity of POT1^35^. The binding of TRF1 to dsTEL might also facilitate and stabilize the binding of POT1 to ssTEL. We also noticed that the 3G-Tract construct showed lower shelterin-bound or POT1-bound states compared to 1G-Tract or 2G-tract constructs. This can be attributed to our recent observation that the 3G-Tract overhang forms a G-Triplex, which inhibits the binding of POT1 and shelterin^36^.

### Protection of the junction region by POT1 and shelterin

Our earlier FRET-PAINT studies on longer telomeric overhangs (up to 24G-Tracts) demonstrated that shelterin reduces accessibility of the overhang more effectively than POT1 alone ^25^. We tested whether the shelterin bridge on telomeric DNA constructs with 1-3 G-tracts in ssTEL and 1-2 repeats in dsTEL can provide effective protection of the junction region. For this, we determined the accessibility of Cy3-labeled 5’-CCC constructs to a Cy5-PNA imager probe that is complementary to ssTEL overhang (Cy5-PNA: 5’-<u>TAACCCT</u>T-Cy5, underlined are complementary to ssTEL) using FRET-PAINT. We observed multiple transient Cy5-PNA binding events in the absence of proteins (Fig. 3B). Regardless of dsTEL length, the frequency of binding events in the DNA-only case increases when the length of ssTEL increases from one to two G-Tracts but remains similar or slightly decreases in the 3G-Tract construct despite having an additional Cy5-PNA binding site (Fig. 3C). This decrease in the binding frequency for 3G-Tract construct can be attributed to G-Triplex formation ^36^, which reduces the accessibility of the overhang to the Cy5-PNA probe.

**Figure 3.**
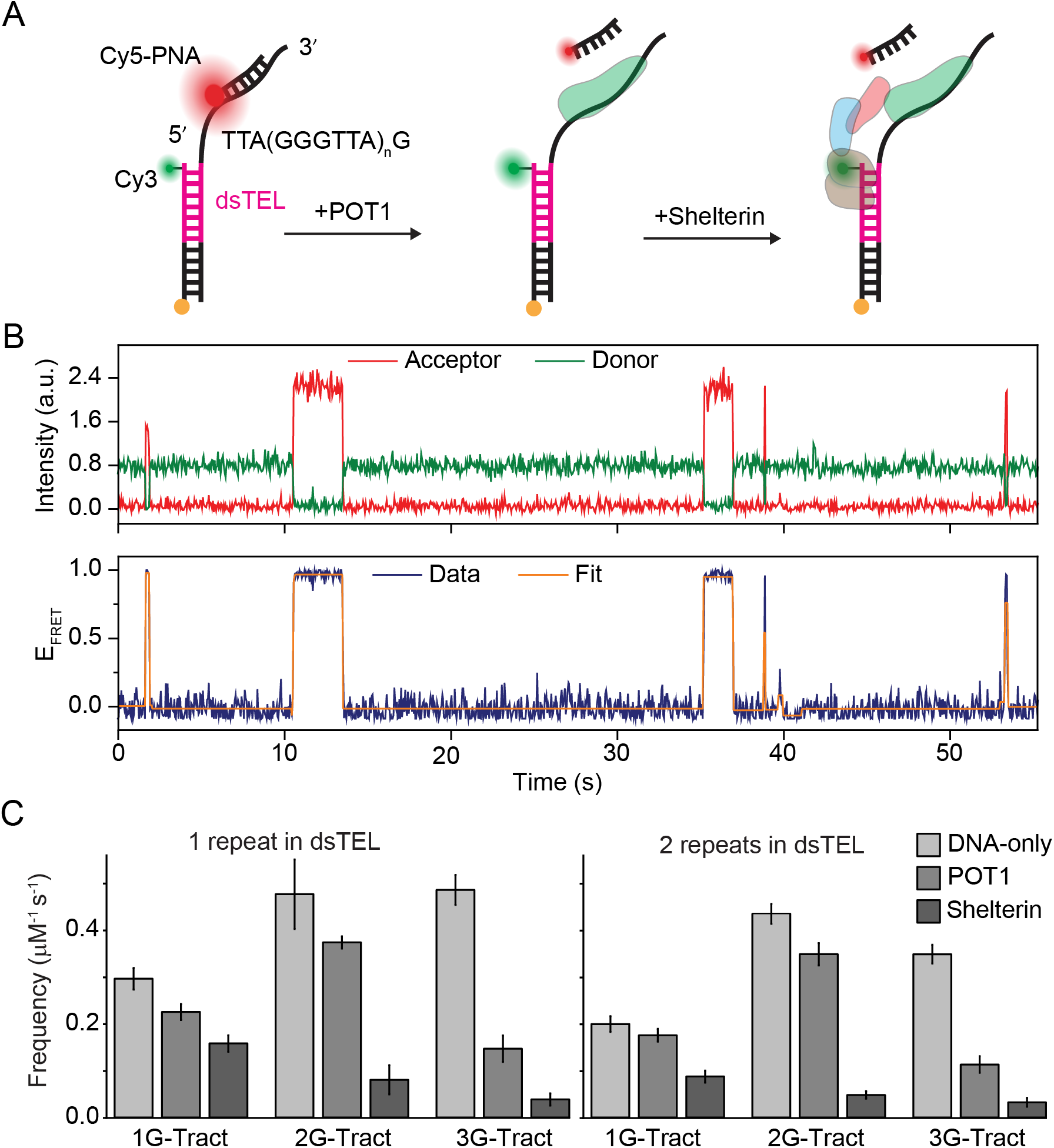
Shelterin sequesters the junction region by forming a bridge between dsTEL and ssTEL. (A) Schematics of 5’-CCC DNA constructs used in FRET-PAINT studies. Cy3 was placed at the junction and Cy5 on the PNA probe which is complementary to the G-rich telomeric sequence. (B) An example time trace showing the FRET bursts due to transient binding of Cy5-PNA to the overhang. (C) Cy5-PNA binding frequencies for constructs containing 1-3 G-Tracts in ssTEL and 1-2 repeats in dsTEL. Error bars were calculated using bootstrapping. Data statistics are given in Table S5 and Table S7. One-way ANOVA analysis showed the variation among different cases for each construct to be statistically significant (p = 0.001, Table S6 and S8).

We anticipated that binding of POT1 or shelterin to the DNA constructs restricts the accessibility of ssTEL, hence reducing the binding frequency of the Cy5-PNA probe. Consistent with this expectation, the accessibility of Cy5-PNA was highest for the DNA-only case for all constructs. To illustrate, in the presence of POT1, the Cy5-PNA binding frequency was 88%, 74%, and 32% of the DNA-only case for 1G-Tract, 2G-Tract, and 3G-Tract overhangs, respectively, for constructs with one repeat in dsTEL. As G-Triplex structures are destabilized by POT1, these structures have a smaller impact on accessibility in the presence of POT1 compared to the DNA-only case. Despite these competing effects, the addition of POT1 resulted in enhanced protection of the 3G-Tract construct, which can be attributed to positive cooperativity of binding of two POT1 molecules to its overhang ^10,35,37^.

In the presence of shelterin, we observed further reduction in the accessibility when the overhang length is increased from 1 to 2 G-Tracts (44% and 11% of the DNA-only case, respectively). However, elongating the overhang to 3-G-Tracts resulted in a modest enhancement (9% of the DNA-only case; Fig. 3C), suggesting that 2 G-Tracts in ssTEL are adequate for effective protection. Similar results were obtained for the shelterin-mediated reduction in Cy5-PNA binding frequency for DNA constructs that contain two dsTEL repeats (Fig. 3C and Table S5-S8). These results suggest a single telomeric repeat in dsTEL is adequate to establish the shelterin bridge, consistent with our observation that the dsTEL repeat closest to the junction is the preferred shelterin binding site (Fig. 1). We also conclude that POT1 alone is not adequate to reduce the accessibility of the junction of 5’-CCC constructs unless multiple POT1 molecules can bind to this region. However, a single POT1 molecule is sufficient to provide effective protection of the junction region when in complex with other shelterin proteins. The level of protection by shelterin is significantly diminished when a single G-Tract is available in ssTEL, suggesting that at least two unfolded G-Tracts are needed for effective binding of POT1 and establishing a stable shelterin bridge.

### Stability of POT1 and shelterin binding to the junction region

To test the binding stability of POT1 and shelterin to the junction region for the 5’-CCC constructs, we performed a protein displacement assay where we challenged these complexes with the unlabeled version of Cy5-PNA (PNA-U). The DNA constructs contained two repeats in dsTEL and were labeled with Cy5 at the 10^th^ bp of dsTEL and with Cy3 at the 3’-end of the overhang for smFRET measurements (Fig. 4A). In the absence of proteins, we observe a single FRET peak for 1G- and 2G-Tract constructs, but two peaks (E_FRET_ = 0.64 ± 0.02 and 0.50 ± 0.02) for the 3G-Tract construct due to folding of three telomeric repeats into the G-Triplex structure ^36^. These two peaks were not distinguishable in Fig. 2B due to the different labeling positions of the FRET dye pairs.

**Figure 4.**
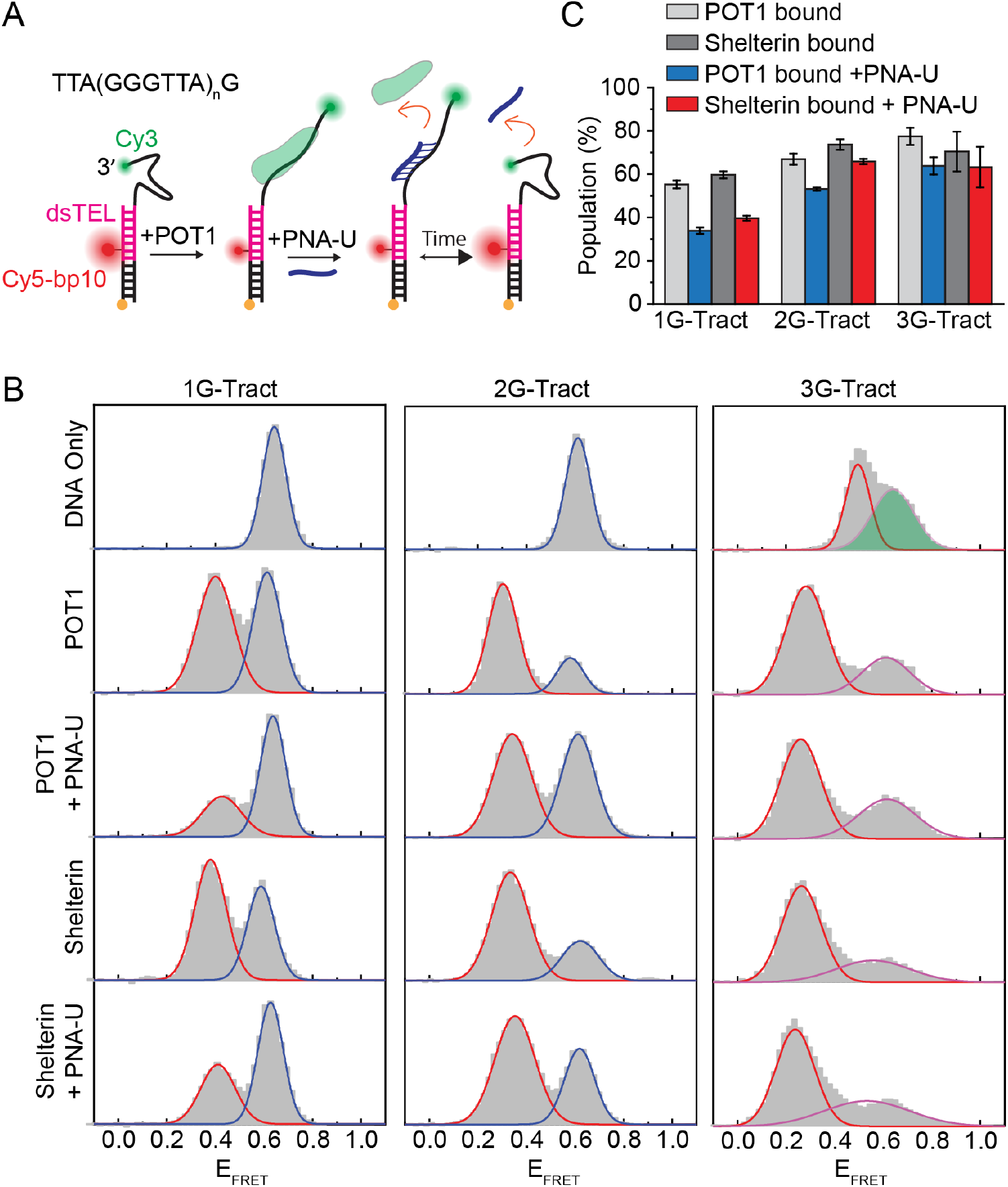
Protein-displacement measurements on 5’-CCC DNA constructs. (A) Schematics of the protein displacement assay. Cy5 is placed at the 10^th^ bp from the junction. Displacement of the bound POT1 or Shelterin from the DNA by PNA-U results in recovering of the higher FRET peak that represents the DNA-only case. (B) Normalized population vs. FRET efficiency histograms for constructs with 1-3 G-Tract overhang (dsTEL contained two repeats). Solid curves represent a fit to a multiple Gaussian. Blue and red curves represent DNA-only and protein-bound populations, respectively. The peak representing G-Triplex formation for the 3G-Tract construct is shaded in green (top, right). The number of molecules and Gaussian fits are given in Table S9 and Table S10. (C) The displacement of POT1 or shelterin by PNA-U is quantified by the areas under fitted Gaussian peaks. The errors are based on fitting errors. T-test analysis showed the variation among different cases for 1G-Tract and 2G-Tract constructs to be statistically significant (p = 0.001, Table S11) while those for 3G-Tract were largely insignificant (p>0.25 for most comparison groups, Table S11).

We incubated the DNA constructs with POT1 or shelterin and determined the fraction of protein-bound DNA molecules from the FRET signal. Then, we introduced PNA-U and determined what fraction of the bound POT1 or shelterin is replaced by PNA-U based on the loss of protein-bound FRET population. Since PNA-U binds only transiently, dissociation of POT1 or shelterin eventually results in the conversion of the lower FRET peak of the protein-bound state to the higher FRET peak of the DNA-only state. 200 nM PNA-U was adequate to displace 21%, 14%, and less than 1% of the bound POT1 molecules from the 1, 2, and 3 G-Tract constructs, respectively. In comparison, PNA-U displaces 20% of shelterin molecules from the 1G-Tract construct, while shelterin displacement was negligible from 2G-Tract and 3G-Tract constructs (Fig. 4C and Table S9-S11). These results agree with our conclusion that POT1 alone requires three G-Tracts in ssTEL to effectively protect 5’-CCC constructs while shelterin achieves that with only two G-Tracts on ssTEL.

### Binding of POT1 and Shelterin to 5’-Phos-CTA Constructs

We also performed measurements on DNA constructs with the CCAATC-5’ sequence in the C-rich strand (5’-CTA), which is identified to be the terminal sequence in the majority of human cells ^30^. In addition, we introduced a phosphate to the 5’-end of this C-rich sequence (5’-Phos-CTA) to investigate its impact on protection by POT1 and shelterin. Due to the modified C-rich strand sequence, all the G-rich overhangs of 5’-Phos-CTA constructs have an additional GG sequence and are two bases longer compared to their 5’-CCC counterparts. This additional GG serves as the fourth G-tract required for GQ formation, resulting in the folding of a 2-tier G-quadruplex in the 3G-Tract constructs of 5’-CTA and 5’-Phos-CTA. This is manifested in the higher E_FRET_ peak for the DNA-only case of 3G-Tract construct compared to the 2G-Tract construct, despite having a longer overhang (Fig. 5). The GQ formation of the 3G-Tract also reduced binding of POT1 and shelterin to the DNA in smFRET measurements (Fig. 5) and lowered the binding frequency of the imager probe to the overhang in FRET-PAINT measurements (Fig. 6).

**Figure 5.**
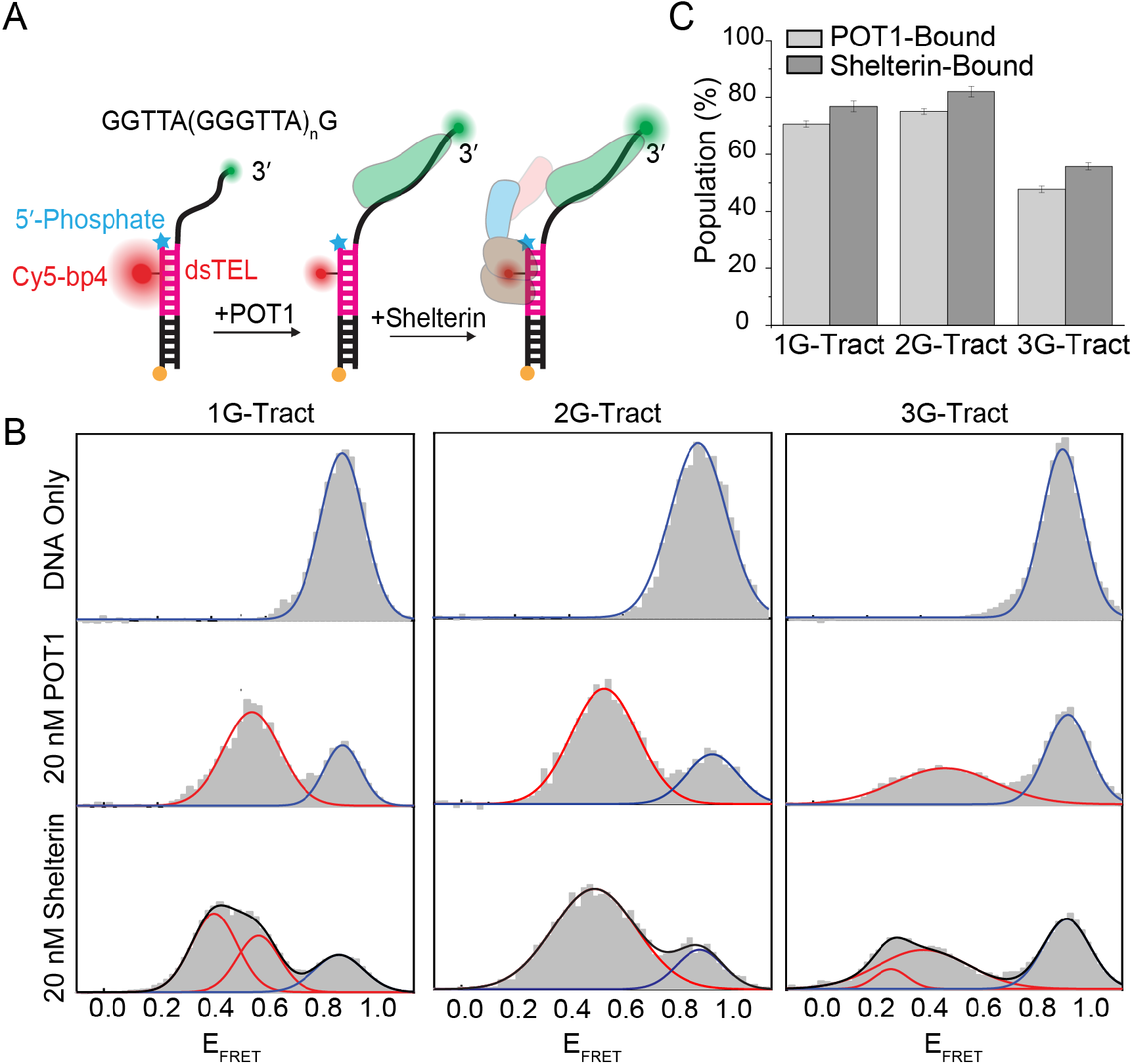
5’-Phos enhances binding efficiency of POT1 to the junction. (A) Schematics of 5’-Phos-CTA DNA constructs in the absence (DNA-only) or the presence of POT1 or shelterin. The C-rich strand terminates with the CCAATC-5’ sequence and contains a phosphate at the 5’-end (cyan star). The Cy5 is placed 4-bp away from the junction and dsTEL contains two repeats. The binding of POT1 or shelterin to the DNA constructs results in emergence of a lower FRET peak. (B) Normalized population vs. FRET efficiency histograms for DNA constructs with 1-3 G-Tracts in ssTEL. Solid curves represent a fit to multiple Gaussian peaks. Blue and red curves represent DNA-only and protein-bound populations, respectively. The black curve (shown only for the shelterin case) is the cumulative of all peaks. The number of molecules for each measurement and the results of Gaussian fits are given in Table S12 and Table S13. (C) Percent population of POT1-bound and shelterin-bound states. In the case of Shelterin-bound states, the total of the two lower FRET peaks (shown in red) was calculated. Error bars were estimated based on fitting errors. T-test analysis showed that the observed differences in populations are statistically significant (p = 0.01-0.03 for all cases, Table S14).

**Figure 6.**
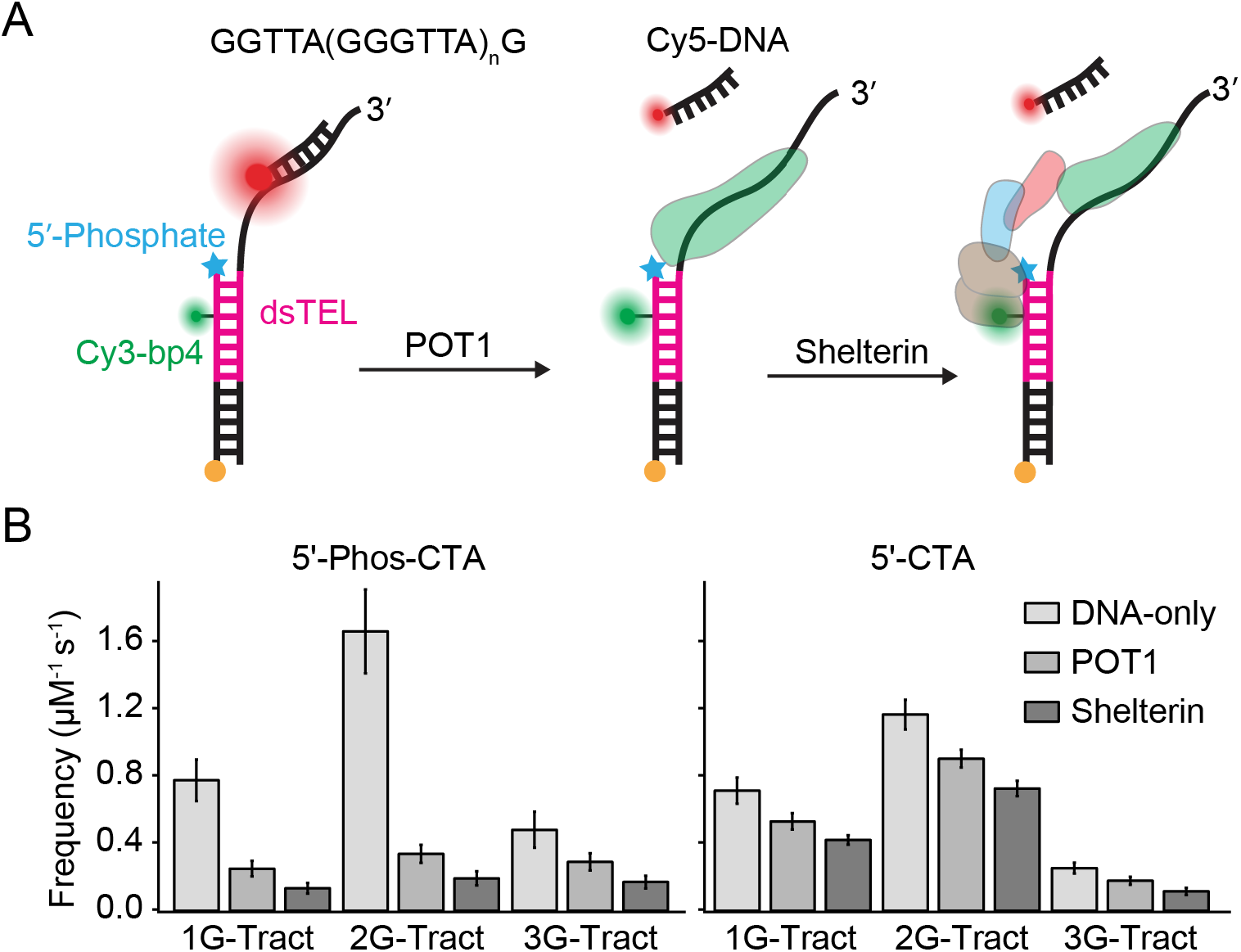
5’-Phos enhances the protection provided by POT1. (A) Schematic of the 5’-Phos-CTA DNA constructs (5’-CTA are identical except the cyan star representing the 5’-phosphate). The C-rich strands of both 5’-CTA and 5’-Phos-CTA constructs end with the CCAATC-5’ sequence, while only the 5’-Phos-CTA constructs have a phosphate modification at the 5’-end. Cy3 is placed 4-bp away from the junction and dsTEL contains two repeats. Cy5-DNA (5’-<u>ACCCTAA</u>A-Cy5) is used as an imager probe in these studies. (B) Cy5-DNA binding frequencies for 5’-Phos-CTA and 5’-CTA constructs, respectively. Error bars are calculated using bootstrapping. Data statistics are given in Table S15 and Table S17. One-way ANOVA test showed that the observed differences in frequencies for each construct are statistically significant (p = 0.001, Table S16 and S18).

POT1 binds very effectively to the 1G-Tract and 2G-Tract 5’-Phos-CTA constructs (Fig. 5, Tables S12-S14). To illustrate, POT1 binds to 71% and 75% of 1G-Tract and 2G-Tract 5’-Phos-CTA constructs compared to 57% and 55% of 5’-CCC constructs (Fig. 2). This is consistent with the recent report that highlights the significance of the 5’-phosphate for recognition of the junction by POT1^24^. Furthermore, the difference in binding affinity of shelterin and POT1, which was consistent in all 5’-CCC constructs (Fig. 2) was less pronounced for 5’-Phos-CTA constructs (Fig. 5). Shelterin was only marginally more efficient than POT1 in binding to 5’-Phos-CTA constructs. The similarity of shelterin and POT1 binding is also extended to the 3G-Tract construct in which destabilization of the GQ is required before POT1 or shelterin can bind to the DNA.

The FRET histograms for the shelterin case show broad distributions that fit well to two Gaussian peaks for 1G-Tract and 3G-Tract constructs, suggesting a heterogeneous distribution of protein-bound states (Fig. 5B). This is particularly unexpected for the 1G-Tract construct, whose overhang is too short to form competing alternative structures or allow binding of multiple POT1 proteins. We propose the lower of these peaks to represent the case where a shelterin bridge is established across the junction whereas the higher peak represents the binding of just the POT1 (TRF1 is bound to dsTEL in the case of lower peak while it is unbound in the other case). In this scenario, when both TRF1 and POT1 are bound across the junction, the ssTEL is stretched more than it would when only POT1 is bound. This is consistent with the shifting of the FRET distributions to lower values in the case of shelterin compared to POT1 alone.

### Impact of 5’-Phosphate on the protection of the junction region by POT1 and Shelterin

For FRET-PAINT studies on 5’-CTA and 5’-Phos-CTA constructs, the donor (Cy3) fluorophore was placed at the 4^th^ nt on the C-rich strand to minimize its impact on binding of proteins to the vicinity of the junction (Fig. 6). The dsTEL contained two repeats in these constructs. Initial tests with these constructs using the Cy5-PNA yielded very few binding events, which we attribute to the modified overhang sequence with an additional GG sequence. We therefore used a modified Cy5-DNA probe (5’-<u>ACC CTA A</u>A-Cy5), which yielded a higher frequency of binding events in the FRET-PAINT studies of both 5’-CTA and 5’-Phos-CTA constructs.

Similar to 5’-CCC constructs, shelterin reduced the accessibility of the overhang more effectively than POT1 alone for all 5’-CTA and 5’-Phos-CTA constructs. However, the impact of POT1 alone was greater in 5’-Phos-CTA compared to those without 5’-phosphate and the difference between POT1 and shelterin-mediated protection was smaller in 5’-Phos-CTA constructs. To illustrate, POT1 binding reduces the Cy5-DNA binding frequency by 66%, 80%, and 40% of the DNA-only case in 1G-Tract, 2G-Tract, and 3G-Tract 5’-Phos-CTA constructs, respectively. The corresponding reductions are 26%, 23%, and 30% in the case of 5’-CTA constructs. These results further highlight the impact of 5’-phosphate on the binding affinity of POT1 to the junction region and suggest less reliance on tethering by shelterin for the physiologically relevant 5’-Phos-CTA constructs. These observations also imply the need for supporting action of shelterin for protection of the junction region when chromosome ends are modified due to nuclease action (as exemplified by the 5-CCC constructs in Figs. 1-4).

Interestingly, the binding frequency of the Cy5-DNA probe was consistently higher in the DNA only cases for 5’-Phos-CTA constructs compared to respective 5’-CTA constructs. To illustrate, the binding frequencies are 12%, 47%, and 95% higher for 1G-Tract, 2G-Tract, and 3G-Tract constructs, respectively, in the 5’-Phos-CTA constructs compared to the respective 5’-CTA constructs. Therefore, even though the 5’-phosphate plays an important role in providing more effective protection by facilitating POT1 binding, it also increases the accessibility of the junction in the absence of POT1. This effect is particularly prominent in the 3G-Tract construct where Cy5-DNA binding frequency increases by 95% in 5’-Phos-CTA constructs compared to the respective 5’-CTA construct. This could be due to the destabilization of the G-quadruplex structure, which inhibits Cy5-DNA binding, by the additional negative charge introduced by the 5’-phosphate.

The junction region is the most accessible region in long telomeric overhangs and the least likely site for GQ formation, which makes protection of this region by POT1 and Shelterin a critical aspect of telomere maintenance. By performing smFRET, FRET-PAINT, PIFE, and protein displacement measurements on minimal telomeric constructs, we showed that a single telomeric repeat on dsTEL (the repeat closest to the junction) and two repeats on ssTEL are adequate for effective protection of the junction region by shelterin in the case of a construct with a TAACCC-5’ sequence at the C-rich strand. Modifying the terminal sequence to CCAATC-5’ and introducing a phosphate to the 5’-end yielded similar results, although POT1-mediated protection was significantly more effective in the presence of the 5’-phosphate. In all studied cases, shelterin reduced ssTEL accessibility more effectively than POT1 alone and enabled the protection of shorter overhangs, further highlighting the significance of forming a shelterin bridge between dsTEL and ssTEL at the junction region.

## Supporting Information

Additional experimental details, materials, and methods, including details of experimental setup and data acquisition parameters, sequences of all DNA constructs and statistical analyses of all presented data.

## Acknowledgements

NIH [1R15GM123443, 1R15GM146180 to H.B.; R35GM136414 to A.Y.] and NSF [MCB 1954449 to A.Y.] funded this work. J.A. is funded by the Deanship of Scientific Research at Northern Border University, Arar, KSA through the project number NBU-SAFIR-2024. Funding for open access charge: NIH Grant and Kent State University Research Office.

## Conflict of interest statement

The authors declare no conflict of interest.

## Notes

### Competing Interest Statement

The authors have declared no competing interest.

