## Supplementary Information for "Protection of the Telomeric Junction by the Shelterin Complex"

### Methods

**DNA Constructs:** Schematics of the assays and partial-duplex DNA (pdDNA) constructs used for smFRET, FRET-PAINT, and PIFE studies are shown in relevant figures. The sequences of all nucleic acids are given in Table S1. The constructs are named based on how many TTAGGG repeats they contain in the overhang and the terminal sequences of the C-rich strands, which dictate the junction sequence. The C-rich strands terminate with either the TAACCC-5′ sequence (called '5′-CCC' constructs to highlight the last three nucleotides) or the CCAATC-5′ sequence (called '5′-CTA' construct). The constructs which have the CCAATC-5′ terminal sequence and a 5′-phosphate at the C-rich stand are identified as 5′-Phos-CTA constructs.

All 5′-CCC constructs contain the number of TTAGGG repeats mention in their name plus a TTAG at the 3′-end, e.g., 2G-Tract construct has the following sequence in the overhang: TTAGGGTTAGGGTTAG-3′. All 5′-CTA and 5′-Phos-CTA constructs contain an additional GG at the 5′-side, e.g., 2G-Tract construct has the following sequence in the overhang: GGTTAGGGTTAGGGTTAG-3′.

The pdDNA constructs were prepared by annealing a long strand, which contains the overhang, and a short strand that includes a random 18-nt sequence and 1-2 repeats of the C-rich telomeric sequence. This C-rich telomeric sequences hybridize with the G-rich telomeric sequences in the long strand to form the dsTEL part of the constructs. The unhybridized G-rich telomeric repeats in the long strand constitute the ssTEL (interchangeably used with 'overhang'). The annealing reaction was performed at 95 °C for 5 minutes and followed by slow cooling to room temperature over 2 hours. For annealing, the short and the long strands were mixed in a molar ratio of 1:4 in a buffer that contains 150 mM KCl and 10 mM $MgCl_2$ (later reduced to 2 mM $MgCl_2$ during smFRET, PIFE, and FRET-PAINT studies).

For the PIFE and smFRET measurements, the Cy5 (acceptor for smFRET) was placed on the short strand while Cy3 (donor for smFRET) was placed at the 3′-end of the long strand (which contains the ssTEL). In the FRET-PAINT measurements, similar pdDNA constructs which are labeled with the donor fluorophore Cy3 at the short strand were utilized. In this case, the acceptor fluorophore is on the imager strand Cy5-PNA (used for 5′-CCC constructs) or Cy5-DNA (used for 5′-CTA and 5′-Phos-CTA constructs). The sequence of Cy5-PNA is 5′-<u>TAACCCT</u>T-Cy5, where underlined are complementary to a 7-nt sequence on ssTEL. The sequence of Cy5-DNA is 5′-<u>ACCCTAA</u>A-Cy5, where underlined are complementary 8-nt sequence on ssTEL.

In order to test for PIFE due to binding of shelterin to different locations on dsTEL, the Cy5 fluorophore was placed at two different sites within C-rich strand of the dsTEL of 5′-CCC constructs: (i) 4 bp away from the junction which is located within the first telomeric repeat (Cy5-bp4); and (ii) 10 bp from the junction which is located within the second telomeric repeat (Cy5-

bp10). Cy5-bp4 construct is expected to yield a PIFE signal if TRF1 binds to the first (from the junction) telomeric repeat on dsTEL, while Cy5-bp10 is expected to show a PIFE signal if TRF1 binds to the second telomeric repeat. These different acceptor positions were also used to optimize the sensitivity of FRET signal to detect binding of POT1 or Shelterin. For consistency of results, PIFE and smFRET measurements were performed on the same constructs within the same sample chambers. Therefore, the Cy3 fluorophore, which is required for smFRET but not PIFE measurements, is kept in the PIFE measurements as well. However, for PIFE measurements the Cy5 fluorophore was directly excited with a λ=632 nm red laser beam (unlike the smFRET measurements in which the Cy3 fluorophore was excited with a λ=532 nm green laser beam), which does not excite Cy3 (<1% of the excitation signal at 532 nm). Therefore, to avoid confusion, we do not show the Cy3 on the PIFE construct in Fig. 1.

**Fluorescence Microscopy Measurements:** A prism-type total internal reflection fluorescence (TIRF) microscope built around an Olympus IX-71 microscope was used for all measurements [27]. Laser drilled quartz slides and glass coverslips were thoroughly cleaned with acetone, 1M potassium hydroxide (KOH), and 20 min of piranha etching, followed by amino silane functionalization for 20-30 min. The slides and coverslips were then passivated with a mixture of mPEG (PEG-5000, Laysan Bio, Inc.) and biotin PEG (biotin-PEG-5000, Laysan Bio, Inc.) in the molar ratio of 40:1 and kept overnight. After cleaning with distilled and deionized water and drying with nitrogen gas, the slides and coverslips were stored at -20 °C. Before the experiments, the slides and coverslips were thawed and passivated with 333 Da PEG for 30-45 min to increase the density of the PEG brush. The microfluidic chambers were created between the slide and coverslip with double-sided tape, followed by sealing the chamber with epoxy. Finally, the microfluidic chamber was treated with 2% (v/v) Tween-20 to reduce non-specific binding. After 5 min incubation, the excess detergent was removed from the chamber.

Before introducing the freshly annealed pdDNA samples (at 10-20 pM concentration in 150 mM KCl and 2 mM $MgCl_2$), 0.01 mg/mL streptavidin was incubated in the chamber for 2 minutes. Excess or unbound DNA were removed from the chamber. All fluorescence measurements were performed in an imaging buffer that contains 50 mM Tris-HCl (pH 7.5), 2 mM Trolox, 0.8 mg/mL glucose, 0.1 mg/mL glucose oxidase, 0.1 mg/mL bovine serum albumin (BSA), 2 mM $MgCl_2$, and 150 mM KCl. For FRET-PAINT studies 40 nM Cy5-PNA or Cy5-DNA was added to the imaging buffer, while for protein displacement assays 200 nM unlabeled PNA (PNA-U) was added to the imaging buffer. In experiments with proteins, 20 nM POT1 or 20 nM shelterin were also included in the imaging buffer. The Cy5-PNA and PNA-U strands were heated to 85 °C for 10 minutes before they were added to the imaging buffer. Short (20 frames) and long movies (1000-2000 frames) were acquired at 100 ms integration time, which were later analyzed using a custom MATLAB code. smFRET histograms were corrected for donor only peak representing $E_{FRET}=0$. In all reported FRET efficiency values, the errors were approximated as the bin size (0.02) of the FRET histogram since the fitting error was smaller than the bin size. Error bars in binding frequencies attained via FRET-PAINT were calculated using the bootstrap method with 2000 sample size as described in an earlier study [20].

**Protein Constructs:** As previously described [28], POT1 and 4-component shelterin (POT1, TPP1, TIN2 and either TRF1 or TRF2) were expressed in insect cells. Briefly, human POT1 with an N terminal ZZ affinity tag, TEV cleavage site, and YBBR labeling site was cloned into an Omnibac vector, and the four-component shelterin containing either TRF1 or TRF2 was cloned into a BigBac vector, where POT1 was given an N terminal YBBR tag, POT1, TRF1 and TRF2 were each given an N terminal ZZ affinity tag and a TEV cleavage site, and TIN2 and TPP1 were each given an N-terminal His-MBP affinity tag and a TEV cleavage site. Protein was purified from insect cells as previously described [29]. Briefly, plasmids containing genes of interest were transformed into DH10Bac competent cells (Berkeley MacroLab), and Bacmid DNA was purified using ZymoPURE miniprep buffers (Zymo Research, D4210) and ethanol precipitation. Insect cells were transfected using Fugene HD transfection reagent (Promega, E2311). The virus was amplified in progressively larger cultures. 1 mL of the P1 virus was used to infect 50 mL of Sf9 cells at 1 million cells/mL for 72 h. 10 mL of the P2 virus was used to infect 1 L of Sf9 cells at 1 million cells/mL and expression proceeded for 72 h. Cells expressing the protein of interest were harvested at 4,000 g for 10 min and resuspended in 50 mL lysis buffer (50 mM HEPES pH 7.4, 1 M NaCl, 10% glycerol, 1 mM PMSF, 1 mM DTT, and 1 tablet of protease inhibitor (Sigma, 4693132001)). Lysis was performed using 15 loose and 15 tight plunges of a Wheaton glass dounce. The lysate was clarified using a 45 min, 360,000 g spin in a Ti70 rotor. The supernatant was incubated with 1 mL IgG beads (IgG Sepharose 6 Fast Flow, GE Healthcare, 17096902) for POT1 or 1 mL amylose beads (New England BioLabs, E8021S) for shelterin for 1 hour. Beads were washed with 40 mL of lysis buffer followed by 40 mL of storage buffer (50 mM Tris pH 7.5, 300 mM KCl, 2 mM $MgCl_2$, 10% glycerol, 1 mM DTT). Beads were then collected and incubated with TEV protease (Berkeley Macrolab, Addgene #8827) for 1 h at room temperature to elute the protein. Cleaved protein was run through a Superdex 200 Increase 10/300 GL size exclusion column (Cytiva, 28-9909-44) to remove TEV, cleaved affinity tags, and shelterin subcomplexes.

### DNA Sequences

The partial duplex DNA (pdDNA) constructs were created by annealing a 30-nt long stem strand with a longer strand that includes a complementary sequence to the stem strand and the telomeric overhang. The following strands were used to create the pdDNA constructs.

| Strands | Sequence in 5′–3′ | Used in Figures |
|---|---|---|
| nG-Tract-Cy3 | TGGCGACGGCAGCGAGGCTTAGGGTTAGGG $(TTAGGG)_n$ TTAG/iCy3 | Figs. 1, 2, 4, 5 |
| nG-Tract | TGGCGACGGCAGCGAGGCTTAGGGTTAGGG $(TTAGGG)_n$ TTAG | Fig. 3, 6 |
| Cy5-bp4 | CCCT /iCy5/AACCCTAA GCCTCGCTGCCGTCGCCA-biotin | Figs. 1, 2 |
| Cy5-bp10 | CCCTAACCCT/iCy5/AA GCCTCGCTGCCGTCGCCA-biotin | Figs. 1, 2, 4 |
| Stem-24 | /Cy3/CCCTAA GCCTCGCTGCCGTCGCCA-biotin | Fig. 3 |
| Stem-30 | /Cy3/CCCTAACCCTAA GCCTCGCTGCCGTCGCCA-biotin | Fig. 3 |
| Phos-Stem-28-Cy5 | Phos-CTAA/iCy5/CCCTAA GCCTCGCTGCCGTCGCCA-biotin | Fig. 5 |
| Stem-28-Cy3 | CTAA/iCy5/CCCTAA GCCTCGCTGCCGTCGCCA-biotin | Fig. 6 |
| Phos-Stem-28-Cy3 | Phos-CTAA/iCy3/CCCTAA GCCTCGCTGCCGTCGCCA-biotin | Fig. 6 |
| Cy5-PNA | TAACCCTT-Cy5 | Fig. 3 |
| PNA-U | TAACCCTT | Fig. 4 |
| Cy5-DNA | ACCCTAAA-Cy5 | Fig. 6 |

**Table S1**: Sequences used for creating pdDNA constructs. The nucleotides in green fonts constitute the overhang for 5'-CCC constructs while the overhangs of 5'-CTA and 5'-Phos-CTA constructs contain an additional GG at the 5'-side. The subscripts designate the number of repeats, i.e. $(GGGTTA)_2$ refers to GGGTTAGGGTTA. Stem strands (24, 28, or 30 nt long) hybridize with a segment of the long strand (24, 28, or 30 nt on the 5'-side) and create a duplex DNA which is attached to the surface via biotin-streptavidin conjugation. The purple nucleotides of the long strands hybridize with the purple nucleotides of the Stem strands and form a 12-bp, 10-bp or 6-bp long telomeric duplex regions for Stem-30, Stem-28, and Stem-24 strands, respectively. In case of Cy5-PNA, PNA-U, and Cy5-DNA strands, the red nucleotides are complementary to the telomeric sequence.

| | Number of Molecules | | |
|---|---|---|---|
| | DNA [illegible] | 20 nM POT1 | 20 nM [illegible] |
| 1G-Tract | 112 | 177 | 228 |
| 2G-Tract | 132 | 257 | 188 |
| 3G-Tract | 83 | 113 | 89 |

**Table S2.** Number of molecules presented in FRET histograms in Figure 2.

| | | DNA Only | | 20 nM POT1 | | 20 nM Shelterin | |
|---|---|---|---|---|---|---|---|
| | | $E_{FRET}$ | Area | $E_{FRET}$ | Area | $E_{FRET}$ | Area |
| 1G-Tract | Peak 2 | | | 0.67 ± 0.02 | 1.18 ±0.01 | 0.66 ±0.02 | 1.57 ± 0.01 |
| | Peak 1 | 0.89±0.02 | 2.09±0.01 | 0.87 ± 0.02 | 0.90 ±0.01 | 0.87 ± 0.02 | 0.55 ± 0.01 |
| 2G-Tract | Peak 2 | | | 0.54 ± 0.02 | 1.23 ±0.02 | 0.50 ± 0.02 | 1.65 ± 0.03 |
| | Peak 1 | 0.86±0.02 | 2.05±0.02 | 0.85 ± 0.02 | 1.01 ±0.02 | 0.82 ± 0.02 | 0.50 ± 0.03 |
| 3G-Tract | Peak 2 | | | 0.42 ± 0.02 | 0.96 ±0.05 | 0.36 ± 0.02 | 1.07 ± 0.06 |
| | Peak 1 | 0.88±0.02 | 2.06±0.04 | 0.89 ± 0.02 | 1.10 ±0.03 | 0.90 ± 0.02 | 0.94 ± 0.04 |

**Table S3:** Gaussian fits for data in Figure 2. Peak 1 represents the DNA-only state. Peak 2 represents the POT1-bound or Shelterin-bound state. The error values associated with the peak positions ($E_{FRET}$) were based on standard error of the peak position. If this error was less than the bin size, the errors were estimated as ±0.02, which is the bin size in the FRET histograms. The populations values reported in the manuscript are based on the relative area under the respective peak. The error values in populations are based on the fitting errors associated with these areas.

| DNA construct Group | t-Test |
|---|---|
| 1G-Tract | t (4) = -11.54, p<0.001 |
| 2G-Tract | t (4) = -18.35, p<0.001 |
| 3G-Tract | t (4) = -5.66, p=0.005 |

**Table S4:** T-Test analysis illustrating the statistical significance of the differences of the peak populations among POT1 and Shelterin cases in Figure 2C.

| | | Frequency ($s^{-1}$) | Total time (s) | Number of binding events | Number of molecules | Error |
|---|---|---|---|---|---|---|
| 1G-Tract | DNA Only | 0.01183 | 21811.43 | 258 | 240 | 0.000926 |
| | POT1 | 0.00899 | 22458.49 | 202 | 190 | 0.000686 |
| | Shelterin | 0.00631 | 19963.02 | 126 | 276 | 0.0007 |
| 2G-Tract | DNA Only | 0.01904 | 25425.84 | 484 | 390 | 0.00296 |
| | POT1 | 0.01493 | 19821.09 | 296 | 401 | 0.000522 |
| | Shelterin | 0.00321 | 21192.34 | 68 | 310 | 0.00124 |
| 3G-Tract | DNA Only | 0.01386 | 27422.49 | 380 | 365 | 0.000802 |
| | POT1 | 0.00449 | 13136.67 | 59 | 280 | 0.00071 |
| | Shelterin | 0.00128 | 12532.23 | 16 | 259 | 0.000378 |

**Table S5:** PNA binding frequencies shown in Figure 3C left panel (one repeat in dsTEL).

| DNA construct Group | ANOVA test |
|---|---|
| 1G-Tract | $F(2,5994) = 26365, p=0.001$ |
| 2G-Tract | $F(2,5994) = 38029, p=0.001$ |
| 3G-Tract | $F(2,5994) = 176413, p=0.001$ |

**Table S6:** One-way ANOVA analysis illustrating the statistical significance of the differences of the frequencies for the DNA-Only, POT1 and Shelterin cases in Fig. 3C left panel (one repeat in dsTEL).

| | | Frequency | Total time | Number of binding events | Number of molecules | Error |
|---|---|---|---|---|---|---|
| 1G-Tract | DNA Only | 0.00798 | 34088.19 | 272 | 360 | 0.000674 |
| | POT1 | 0.00702 | 33182.4 | 233 | 350 | 0.000549 |
| | Shelterin | 0.00349 | 19492.88 | 68 | 200 | 0.000503 |
| 2G-Tract | DNA Only | 0.0174 | 37524.66 | 653 | 320 | 0.000864 |
| | POT1 | 0.01394 | 28759.56 | 401 | 255 | 0.000958 |
| | Shelterin | 0.00192 | 40217.72 | 77 | 366 | 0.000318 |
| 3G-Tract | DNA Only | 0.01386 | 27422.49 | 380 | 365 | 0.000802 |
| | POT1 | 0.00449 | 13136.67 | 59 | 280 | 0.00071 |
| | Shelterin | 0.00128 | 12532.23 | 16 | 259 | 0.000378 |

**Table S7:** PNA binding frequencies shown in Figure 3C right panel (two repeats in dsTEL).

| DNA construct Group | ANOVA test |
|---|---|
| 1G-Tract | F (2,5994) = 36565, p=0.001 |
| 2G-Tract | F (2,5994) = 261017, p=0.001 |
| 3G-Tract | F (2,5994) = 206441, p=0.001 |

**Table S8:** One-way ANOVA analysis illustrating the statistical significance of the differences of the frequencies for the DNA-Only, POT1 and Shelterin cases in Fig. 3C right panel (two repeats in dsTEL).

| | Number of Molecules | | | | |
|---|---|---|---|---|---|
| | DNA Only | 20 nM POT1 | 200 nM PAN-U | 20 nM Shelterin | 200 nM PAN-U |
| 1G-Tract | 115 | 160 | 140 | 110 | 123 |
| 2G-Tract | 153 | 215 | 184 | 105 | 151 |
| 3G-Tract | 208 | 128 | 131 | 160 | 116 |

**Table S9.** Number of molecules presented in FRET histograms in Figure 4.

| | | 1G-Tract | 2G-Tract | 3G-Tract |
|---|---|---|---|---|
| DNA Only | Peak $E_{FRET}$ | 0.64 ± 0.02 | 0.61 ± 0.02 | 0.50 ± 0.02<br>0.64 ± 0.02 |
| | Area | 2.13 ± 0.02 | 2.04 ± 0.02 | 0.97 ± 0.04<br>1.18 ± 0.04 |
| 20 nM POT1 | Peak $E_{FRET}$ | 0.40 ± 0.02<br>0.61 ± 0.02 | 0.31 ± 0.02<br>0.58 ± 0.02 | 0.30 ± 0.02<br>0.58 ± 0.02 |
| | Area | 1.21 ± 0.02<br>0.98 ± 0.02 | 1.43 ± 0.02<br>0.71 ± 0.02 | 1.65 ± 0.02<br>0.48 ± 0.02 |
| 200 nM PNA-U after POT1 | Peak $E_{FRET}$ | 0.43 ± 0.02<br>0.64 ± 0.02 | 0.34 ± 0.02<br>0.61 ± 0.02 | 0.26 ± 0.02<br>0.62 ± 0.02 |
| | Area | 0.73 ± 0.02<br>1.43 ± 0.02 | 1.13 ± 0.01<br>1.00 ± 0.01 | 1.42 ± 0.03<br>0.81 ± 0.04 |
| 20 nM Shelterin | Peak $E_{FRET}$ | 0.38 ± 0.02<br>0.59 ± 0.02 | 0.33 ± 0.02<br>0.62 ± 0.02 | 0.26 ± 0.02<br>0.56 ± 0.05 |
| | Area | 1.26 ± 0.02<br>0.85 ± 0.01 | 1.62 ± 0.01<br>0.58 ± 0.01 | 1.50 ± 0.05<br>0.63 ± 0.06 |
| 200 nM PNA-U after Shelterin | Peak $E_{FRET}$ | 0.41 ± 0.02<br>0.63 ± 0.02 | 0.35 ± 0.02<br>0.62 ± 0.02 | 0.35 ± 0.02<br>0.62 ± 0.02 |
| | Area | 0.85 ± 0.02<br>1.29 ± 0.01 | 1.41 ± 0.01<br>0.73 ± 0.01 | 1.41 ± 0.01<br>0.73 ± 0.01 |

**Table S10:** Gaussian fits for data in Figure 4. The error values associated with each peak were based on standard error of the peak position. If this error was less than the bin size, the errors were estimated as ±0.02, which is the bin size in the histograms. The populations values reported in the manuscript are based on the relative area under the respective peak. The error values in populations are based on the fitting errors associated with these areas.

| DNA construct group | | t-Test |
|---|---|---|
| 1G-Tract | POT1 *vs* POT1+PNA-U | t (4) = 13.29, p<0.001 |
| | POT1 *vs* Shelterin | t (4) = -4.09, p=0.015 |
| | POT1+PNA-U *vs* Shelterin+PNA-U | t (4) = -6.39, p=0.003 |
| | Shelterin *vs* Shelterin+PNA-U | t (4) = 15.59, p<0.001 |
| 2G-Tract | POT1 *vs* POT1+PNA-U | t (4) = 4.90, p=0.008 |
| | POT1 *vs* Shelterin | t (4) = -2.86, p=0.046 |
| | POT1+PNA-U *vs* Shelterin+PNA-U | t (4) = -3.36, p=0.028 |
| | Shelterin *vs* Shelterin+PNA-U | t (4) = 3.54, p=0.024 |
| 3G-Tract | POT1 *vs* POT1+PNA-U | t (4) = 3.33, p=0.029 |
| | POT1 *vs* Shelterin | t (4) = 0.43, p=0.692 |
| | POT1+PNA-U *vs* Shelterin+PNA-U | t (4) = -0.52, p=0.63 |
| | Shelterin *vs* Shelterin+PNA-U | t (4) = 1.34, p=0.251 |

**Table S11:** T-Test analysis illustrating the statistical significance of the differences of the peak populations among POT1 and Shelterin cases in Figure 4C. The differences for comparisons within 1G-Tract and 2G-Tract groups are statistically significant; however, those within 3G-Tract group (except for POT1 vs POT1+PNA-U comparison) are not (p=0.25-0.69).

| | Number of Molecules | | |
|---|---|---|---|
| | DNA Only | 20 nM POT1 | 20 nM Shelterin |
| 1G-Tract | 113 | 143 | 226 |
| 2G-Tract | 123 | 139 | 158 |
| 3G-Tract | 165 | 189 | 195 |

**Table S12.** Number of molecules presented in FRET histograms in Figure 5.

| | | DNA Only | | 20 nM POT1 | | 20 nM Shelterin | |
|---|---|---|---|---|---|---|---|
| | | EFRET | Area | EFRET | Area | EFRET | Area |
| 1G-Tract | Peak 3 | | | | | 0.41 ±0.02 | 0.99 ± 0.12 |
| | Peak 2 | | | 0.55 ± 0.02 | 1.45 ±0.04 | 0.57 ±0.02 | 0.65 ± 0.12 |
| | Peak 1 | 0.88±0.02 | 1.99±0.02 | 0.89 ± 0.02 | 0.60 ±0.03 | 0.87 ± 0.02 | 0.49 ± 0.02 |
| 2G-Tract | Peak 3 | | | | | | |
| | Peak 2 | | | 0.53 ± 0.02 | 1.63 ±0.03 | 0.49 ± 0.02 | 1.79 ± 0.07 |
| | Peak 1 | 0.89±0.02 | 2.10±0.02 | 0.93 ± 0.02 | 0.54 ± 0.03 | 0.89 ± 0.02 | 0.39 ± 0.03 |
| 3G-Tract | Peak 3 | | | | | 0.29 ± 0.02 | 0.18 ± 0.05 |
| | Peak 2 | | | 0.49 ± 0.02 | 0.99 ±0.04 | 0.41 ±0.02 | 0.97 ±0.07 |
| | Peak 1 | 0.93±0.02 | 1.91±0.03 | 0.94 ± 0.02 | 1.08 ±0.02 | 0.94 ± 0.02 | 0.91 ± 0.02 |

**Table S13:** Gaussian fits for data in Figure 5. Peak 1 represents the DNA-only state. Peak 2 under 20 nM POT1 represents the POT1-bound state. Peak 2 and 3 under 20 nM Shelterin represent the Shelterin-bound states. The error values associated with the peak positions ($E_{FRET}$) were based on the standard error of the peak position. If this error was less than the bin size, the errors were estimated as ±0.02, which is the bin size in the FRET histograms. The population values reported in the manuscript are based on the relative area under the respective peak. The error values in populations are based on the fitting errors associated with these areas.

| DNA construct group | T-Test |
|---|---|
| 1G-Tract | t (4) = -3.28, p=0.03 |
| 2G-Tract | t (4) = -3.60, p=0.02 |
| 3G-Tract | t (4) = -5.40, p=0.01 |

**Table S14:** T-test analysis illustrating the statistical significance of the differences of the peak populations among POT1 and Shelterin cases in Figure 5C.

| | | Frequency ($s^{-1}$) | Total time (s) | Number of binding events | Number of molecules | Error |
|---|---|---|---|---|---|---|
| 1G-Tract | DNA Only | 0.03112 | 7936.94 | 247 | 309 | 0.004961 |
| | POT1 | 0.01005 | 11348.18 | 114 | 402 | 0.001860 |
| | Shelterin | 0.00537 | 12473.73 | 67 | 542 | 0.001271 |
| 2G-Tract | DNA Only | 0.06661 | 5344.40 | 356 | 161 | 0.249766 |
| | POT1 | 0.01358 | 11047.84 | 150 | 435 | 0.053367 |
| | Shelterin | 0.00774 | 11113.98 | 86 | 443 | 0.041978 |
| 3G-Tract | DNA Only | 0.01931 | 9580.96 | 185 | 346 | 0.107447 |
| | POT1 | 0.01167 | 7882.71 | 92 | 309 | 0.051322 |
| | Shelterin | 0.00686 | 9618.50 | 66 | 398 | 0.038159 |

**Table S15:** DNA binding frequencies for 5'-Phos-CTA Constructs in Figure 6B.

| DNA construct Group (with 5'-Phosphate) | ANOVA test |
|---|---|
| 1G-Tract | $F(2,594) = 4757$, $p=0.001$ |
| 2G-Tract | $F(2,594) = 7480$, $p=0.001$ |
| 3G-Tract | $F(2,594) = 803$, $p=0.001$ |

**Table S16:** One-way ANOVA analysis illustrating the statistical significance of the differences of the frequencies for the DNA-Only, POT1 and Shelterin cases for 5'-Phos-CTA Constructs in Fig. 6B.

| | | Frequency ($s^{-1}$) | Total time (s) | Number of binding events | Number of molecules | Error |
|---|---|---|---|---|---|---|
| 1G-Tract | DNA Only | 0.02776 | 15420.63 | 428 | 265 | 0.003016 |
| | POT1 | 0.02065 | 12980.70 | 268 | 186 | 0.001889 |
| | Shelterin | 0.01635 | 8380.22 | 137 | 143 | 0.001082 |
| 2G-Tract | DNA Only | 0.04532 | 4478.80 | 203 | 212 | 0.003434 |
| | POT1 | 0.03513 | 6206.04 | 218 | 238 | 0.002061 |
| | Shelterin | 0.02824 | 5808.03 | 164 | 214 | 0.001796 |
| 3G-Tract | DNA Only | 0.00988 | 24601.34 | 243 | 466 | 0.001246 |
| | POT1 | 0.00696 | 11067.18 | 77 | 252 | 0.000917 |
| | Shelterin | 0.00448 | 8696.53 | 39 | 237 | 0.000826 |

**Table S17:** DNA binding frequencies for 5′-CTA constructs in Figure 6B.

| DNA construct Group (without 5'-Phosphate) | ANOVA test |
|---|---|
| 1G-Tract | $F(2,594) = 682$, $p=0.001$ |
| 2G-Tract | $F(2,594) = 690$, $p=0.001$ |
| 3G-Tract | $F(2,594) = 602$, $p=0.001$ |

**Table S18:** One-way ANOVA analysis illustrating the statistical significance of the differences of the frequencies for the DNA-Only, POT1 and Shelterin cases for 5′-CTA constructs in Fig. 6B.
